# A Brugada-related KCNT1 mutation unveils its conductance-independent activation of store-operated Ca^2+^ entry

**DOI:** 10.64898/2026.06.12.731813

**Authors:** Pei-Ju Tsai, Yu-Chiao Lin, Hsuan-Chao Lin, You-Yi Li, Jyh-Ming Jimmy Juang, Chien-Yuan Pan, Wen-Pin Chen, Hsi-Chun Chao, Feng-Chiao Tsai

## Abstract

**Background:** Potassium sodium-activated channel subfamily T member 1 (KCNT1) is a Na^+^-activated K^+^ channel, associated with epilepsy and cardiac diseases. Besides K^+^ conductance, KCNT1 is also involved in Ca^2+^ handling. *KCNT1:R1106Q* was identified in a Brugada Syndrome (BrS) patient; however, its pathological characteristics regarding either K^+^ or Ca^2+^ homeostasis remain to be fully elucidated.

**Methods:** Fura-2-loaded HEK293T cells were used for intracellular Ca^2+^ investigations, and whole-cell patch-clamp technique were used as complementary tests. Stromal interaction molecule 1 (STIM1) aggregation and endoplasmic reticulum (ER)-plasma membrane (PM) junctions in HeLa cells were visualized using spinning disc confocal microscope. Multielectrode array (MEA) assays were utilized to assess the field potential duration (FPD) and spontaneous beating rate of human induced pluripotent stem cell-derived cardiomyocytes (hiPSC-CMs).

**Results:** KCNT1 overexpression increased SOCE, which was further upregulated by the KCNT1^R1106Q^ mutant. Furthermore, KCNT1 upregulated ER Ca^2+^ release and was found to localize at ER-PM junctions. The ER Ca^2+^ dynamics in KCNT1^R1106^–overexpressing cells was disrupted, while its SOCE remained intact. The KCNT1 cytoplasmic tail alone sufficiently modulated Ca^2+^ homeostasis. Given that KCNT1 possesses prominent charge-concentrated segments within its cytoplasmic domain, we substituted charged amino acids in either the 740-DDE-742 or 1114-RRLSR-1118 motifs with alanine; these substitutions abolished the KCNT1-mediated increases in ER Ca^2+^ release and ER-PM contact formation. MEA recordings of hiPSC-CMs revealed that the overexpression of KCNT1 shortened the FPD and accelerated the cardiomyocyte beating rate compared to baseline. However, the magnitude of beating rate acceleration induced by KCNT1^R1106Q^ was significantly attenuated compared to that of KCNT1^WT^.

**Conclusions:** Our findings indicate that, independent of KCNT1-mediated ion conductance, the charged motifs located at both ends of the KCNT1 cytoplasmic tail serve as structural anchors to facilitate ER-PM contact formation. This non-conducting role endows KCNT1 with a K^+^ current-independent mechanism to modulate intracellular Ca^2+^ homeostasis and cardiomyocyte physiology.

**Graphical abstract:** 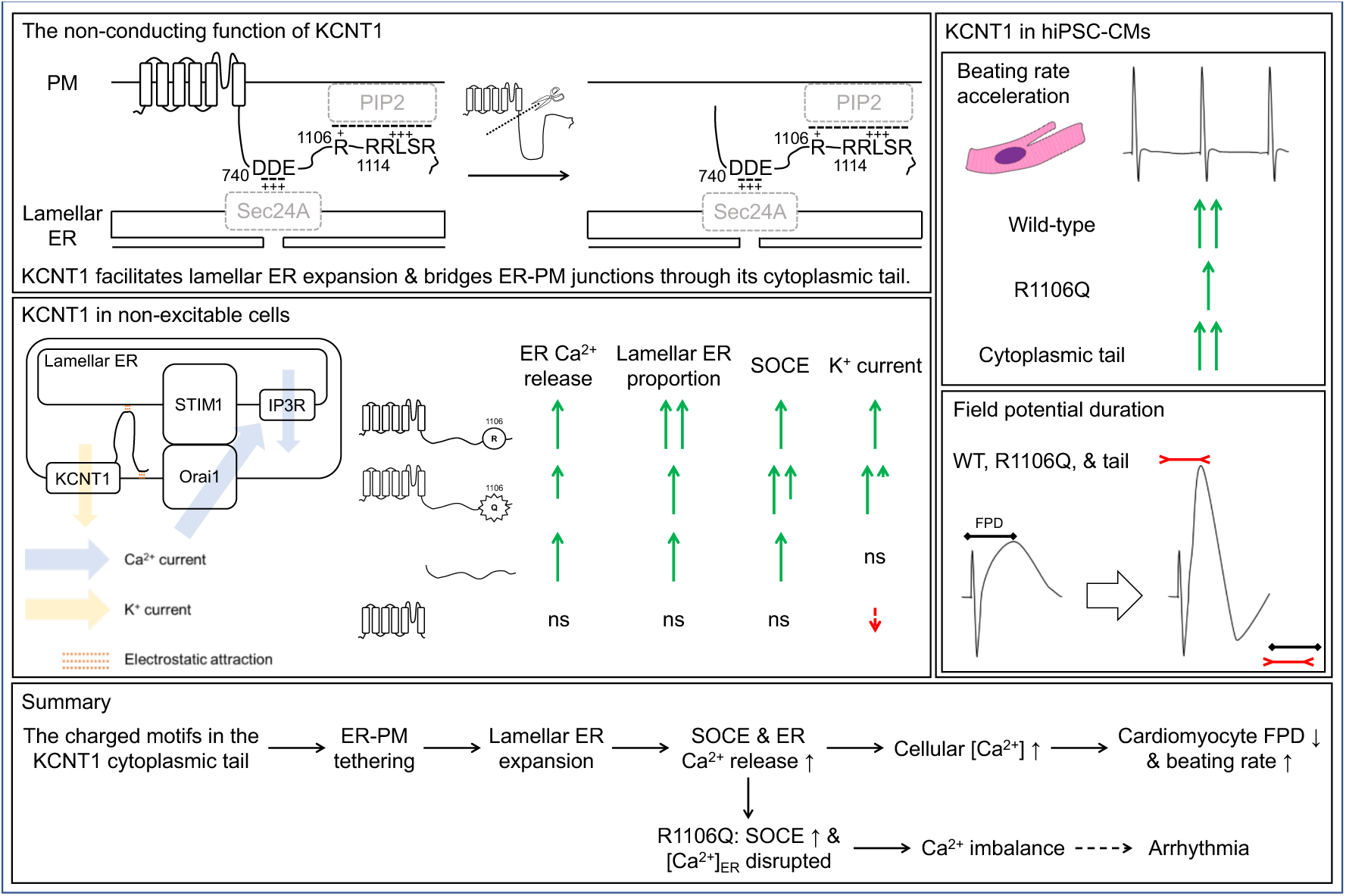

**Novelty and Significance:** *What is known?:*

1. KCNT1 is a Na^+^-activated K^+^ channel, and mutations within this gene are often associated with epilepsy and epilepsy-related cardiac diseases.
2. Besides functioning as a rectifier, KCNT1 is implicated in cellular Ca^2+^ handling, such as regulating Ca^2+^ oscillations in rat cortical neurons.
3. *KCNT1:R1106Q* missense mutation has been identified in a patient with Brugada Syndrome (BrS); however, its pathophysiological characteristics and underlying mechanisms remain uncharacterized.

**What New Information Does This Article Contribute?:**

1. KCNT1 overexpression upregulates both ER Ca^2+^ release and store-operated Ca^2+^ entry (SOCE). KCNT1^R1106Q^ further increased SOCE intensity, however, its increase in ER Ca^2+^ release and [Ca^2+^]_ER_ were attenuated.
2. The cytoplasmic tail of KCNT1 alone is suNicient to modulate Ca^2+^ closely mimicking the eNects of full-length KCNT1.
3. The charged clusters distributed along the KCNT1 cytoplasmic tail facilitate ER-PM contact. Alanine substitutions abolish the enhancements in ER Ca^2+^ release and ER-PM contact formation, while SOCE modulation remains intact.
4. The KCNT1 cytoplasmic tail alone shortens the field potential duration (FPD) and accelerates the spontaneous beating rate in hiPSC-CMs. However, the magnitude of beating rate acceleration induced by KCNT1^R1106Q^ is significantly attenuated compared to KCNT1^WT^.

This study uncovers a novel, conductance-independent role for the KCNT1 channel in structural organization, demonstrating that the charged motifs on its cytoplasmic tail serve as physical anchors facilitating ER-PM junctions. This structural mechanism regulates intracellular Ca2+ homeostasis and subsequently alters cardiomyocyte electrophysiology. By decoupling KCNT1’s structural impact from its ion-conducting function, these findings provide new mechanistic insights into how specific KCNT1 variants, such as the BrS-associated KCNT1:R1106Q mutation, may contribute to arrhythmogenesis through imbalanced Ca2+ handling rather than classical K+ current alterations.

## Introduction

BrS is an inherited, life-threatening heart rhythm disorder that causes sudden death due to ventricular fibrillation. It has a worldwide pooled prevalence of 0.5 per 1000, peaking at 3.7 per 1000 in Southeast Asia ^1–3^. The typical electrocardiography (ECG) of BrS patients exhibits characteristically elevated ST segments in the right precordial leads (V1–V3) ^4,5^.

While there are reports about BrS phenocopies induced by environmental factors such as body temperature, electrolyte imbalance, and certain medications, ion channelopathies leading to abnormal ion current density or alteration in channel expression level are often been identified ^6,7^. Several BrS-related Na^+^, Ca^2+^, and K^+^ channel mutations have been documented; mutations in the former two are generally classified as *loss-of-function* (*LOF*), whereas mutations in the latter are categorized as *gain-of-function* (*GOF*) ^8^.

In 2014, a point mutation in *KCNT1* (*KCNT1:R1106Q*) was identified in a BrS patient through disease-targeted multi-gene sequencing ^9^. KCNT1 is predominantly expressed in the central nervous system (CNS) and heart. To date, most *KCNT1* mutations have been associated with epilepsy and intellectual disability ^10–12^. Besides the CNS, KCNT1 has also been implicated in cardioprotective signaling, cardiac arrhythmia, and blood pressure moderation ^13–15^.

Although the *KCNT1:R1106Q* mutation has been identified, its pathophysiological impact on KCNT1 characteristics remains unclear. Notably, a previous study reported that a patient with epilepsy of infancy with migrating focal seizures (EIMFS) carrying a different missense mutation at the same residue (*KCNT1:R1106P*) did not exhibit cardiac rhythm abnormalities ^16^.

As a Na^+^-activated K^+^ channel that participates in the late afterhyperpolarization response to repetitive firing ^17,18^, KCNT1 consists of N-terminus that regulates its trafficking and heteromeric tetramer formation, six transmembrane domains, and a bulky cytoplasmic tail. The cytoplasmic tail can be subdivided into two consecutive regulators of K^+^ conductance, RCK1 and RCK2. Within the RCK2 domain, a nicotinamide adenine dinucleotide (NAD^+^) binding motif facilitates Na^+^ binding and channel sensitization ^19–21^.

Both *LOF* and *GOF* mutations can lead to cellular hyperexcitability. While it is more intuitively straightforward that *LOF* mutations of K^+^ channels promote excitatory postsynaptic potentials (EPSPs) and increase action potential (AP) firing frequency, linking K^+^ channel *GOF* mutation with epilepsy appears counterintuitive ^22^. However, research on the autosomal dominant nocturnal frontal lobe epilepsy (ADNFLE)-associated *KCNT1:Y796H* mutation demonstrated that such mutations can exert neuron-subtype-specific effects in GABAergic neurons, resulting in network hyperexcitability, and seizures ^23^.

In addition to modulating K^+^ current, Ca^2+^ activities were also affected by KCNT1. The inhibition of KCNT1 has been shown to decrease the frequency of baseline intracellular Ca^2+^ oscillations in cultured rat cortical neurons ^24^. Furthermore, GCaMP6s-expressing transgenic mice have been utilized to map cortical regions impacted by *Kcnt1* variants ^23^. More broadly, inhibiting Kv channels in rat cortical astrocytes suppresses SOCE ^25^, a crucial mechanism for replenishing intracellular Ca^2+^ store following ER Ca^2+^ exhaustion ^26^.

Given that BrS-associated K^+^ channel mutations are classified as *GOF*, and that KCNT1 is involved in various Ca^2+^ activities, we hypothesized that *KCNT1:R1106Q* is a *GOF* mutation which exaggerates KCNT1’s influence on Ca^2+^ homeostasis. In this study, we found that KCNT1 upregulated not only SOCE intensity, but also the amount of Ca^2+^ released from the ER. Moreover, we demonstrate that expressing the cytoplasmic tail of KCNT1 is sufficient to alter Ca^2+^ homeostasis, shorten the FPD in hiPSC-CMs, and accelerate the spontaneous beating rate. Conversely, truncating the cytoplasmic tail abolishes these changes in SOCE and ER Ca^2+^ release.

Furthermore, substituting charge-carrying amino acids in either the 740-DDE-742 or 1114-RRLSR-1118 motifs reversed the increased ER Ca^2+^ release and reduced the ER-PM contact area, implicating the KCNT1 cytoplasmic tail as a crucial physical bridge between the ER and the PM. Our results reveal a novel, K^+^ conductance-independent mechanism by which KCNT1 modulates Ca^2+^ homeostasis, thereby altering cardiomyocyte physiology.

## Results

### KCNT1 localizes at ER-PM junctions and enhances cellular both ER Ca^2+^ release and SOCE

Given that several BrS-related K^+^ channel mutations are classified *GOF* variants and that Kv channel activities can modulate SOCE, we hypothesized that KCNT1^R1106Q^ would similarly enhance Ca^2+^ flow. To evaluate whether KCNT1 alters Ca^2+^ homeostasis, we transfected HEK293T cells with KCNT1 and loaded with Fura-2/AM to monitor changes in the 340/380 nm fluorescence ratio.

The peak Fura-2 ratio following thapsigargin and EGTA treatment was used to quantify ER Ca^2+^ release, while the peak following CaCl_2_ addition represented SOCE intensity (Figure 1A, B) ^27^. ER Ca^2+^ release was significantly increased in KCNT1-overexpressing cells, while there was no statistically significant difference between KCNT1^WT^ and KCNT1^R1106Q^, all of the data points in the KCNT1^R1106Q^ group were actually lower than the average of KCNT1^WT^ group (Figure 1B-D). Using T1ER, a FRET-based ER Ca^2+^ indicator ^28^, the relative ER Ca^2+^ concentration ([Ca^2+^]_ER_) was enhanced by KCNT1^WT^ overexpression, and the FRET ratio of the KCNT1^R1106Q^ group was only slightly higher than the mCherry control (Figure S1A, B). As expected, KCNT1 overexpression led to elevated SOCE, and further potentiated by the *KCNT1:R1106Q* mutation (Figure 1B, C, E). To verify the source of this upregulated signal, we applied BTP2, a potent SOCE inhibitor ^29^, which abolished most of the CaCl_2_-induced Ca^2+^ influx (Figure 1F, G). Furthermore, this BTP2-sensitive Ca^2+^ current was downregulated upon the knockdown of either STIM1 or Orai1, both of which are essential components of the SOCE complex (Figure S1C, D) ^26^.

**Figure 1.**
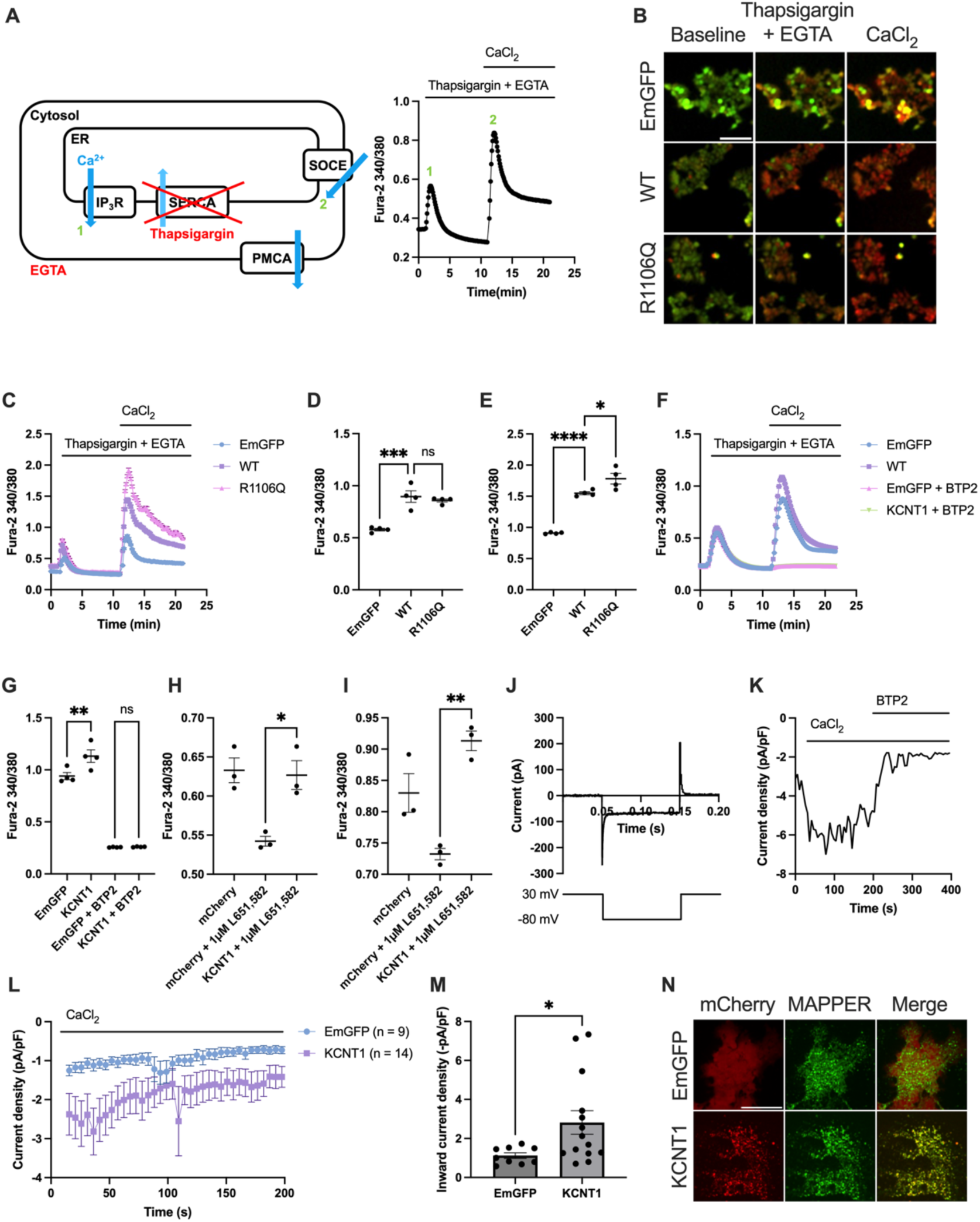
KCNT1 increases ER Ca^2+^ release and SOCE, and localizes to ER-PM junction. (A) Schematic representation of the SOCE assay. The amplitude of ER Ca^2+^ release (1) was evaluated as the peak of Fura-2 ratio following the application of thapsigargin and EGTA. SOCE (2) was measured as the peak ratio following CaCl_2_ addition. (B) Representative ratiometric images of Fura-2 340 nm (red) / 380 nm (green) fluorescence during the SOCE assay. Scale bar: 100 µm. (C) Representative traces of Fura-2 340/380 ratio changes in HEK293T cells. (D) Both KCNT1^WT^ and KCNT1^R1106Q^ significantly elevated ER Ca^2+^ release compared to the EmGFP control. However, all the data points of KCNT1^R1106Q^ were lower than the average of KCNT1^WT^ group. (E) KCNT1^WT^ increased SOCE, which was further amplified by the *KCNT1:R1106Q* mutation. (F) Fura-2 ratio traces in the presence of the SOCE inhibitor BTP2. (G) BTP2 effectively suppressed CaCl_2_-induced Ca^2+^ influx in ER Ca^2+^-depleted HEK293T cells. (H-I) Quantification of ER Ca^2+^ release (H) and SOCE intensity (I) in the presence of 1 µM L-651,582. Although overall signals were suppressed, KCNT1 overexpression consistently upregulated both parameters. (J) Voltage protocol for ICRAC measurement. Cells were held at +30 mV, and I_CRAC_ was recorded during 100-ms voltage steps to −80 mV. (K) Representative recording of I_CRAC_ amplitude in an ER Ca^2+^-depleted HEK293T cell, demonstrating its responses to CaCl_2_ application and subsequent BTP2 inhibition. (L) Time course of I_CRAC_ development following CaCl_2_ administration in ER Ca^2+^-depleted HEK293T cells. (M) Quantification of peak inward current density. KCNT1-expressing cells exhibited significantly larger inward currents than EmGFP controls. (N) Representative images at the basal focal plane of HeLa cells. The punctate distribution of KCNT1 colocalized with the ER-PM junction marker MAPPER. Scale bar: 20 µm. \**p < 0.05*. Image contrast in (N) was adjusted independently.

Since Ca^2+^ influx through L-type Ca^2+^ channels (LTCC) is critical for the cardiac AP plateau ^30^. We sought to exclude LTCC interference by treating cells with 1 or 10 µM L-651,582, a L-type voltage-dependent Ca^2+^ channel (VDCC) blocker ^31^, during the Fura-2 loading period. Although 1 µM L-651,582 suppressed overall ER Ca^2+^ release and SOCE, KCNT1 overexpression still significantly upregulated both parameters compared to the mCherry control (Figure S1E, Figure 1H, I). Even under the substantial disruption of Ca^2+^ homeostasis caused by 10 µM L-651,582, KCNT1-overexpressing cells maintained a trend toward increased ER Ca^2+^ release and SOCE (Figure S1F-H).

Given KCNT1’s abundance in the CNS, we also performed SOCE assay in SH-SY5Y cells, which exhibit significantly higher endogenous KCNT1 levels than HEK293T cells (Figure S1I). Similar our findings in HEK293T cells, KCNT1-overexpression in SH-SY5Y cells increased both ER Ca^2+^ release and SOCE intensity (Figure S1J, K). Conversely, KCNT1 knockdown SH-SY5Y cells led to a significant reduction in SOCE, yet the ER Ca^2+^ release did not show a significant decrease (Figure S1L, M).

To further validate these findings, we measured I_CRAC_ using whole-cell patch-clamp recording, a reliable method for exploring inward current following ER Ca^2+^ depletion ^32^. KCNT1 significantly amplified I_CRAC_ significantly upon the introduction of CaCl_2_ in tetrodotoxin-pretreated, ER Ca^2+^ depleted cells, consistent with our Fura-2 imaging results (Figure 1J-M).

Because KCNT1 modulated both ER Ca^2+^ release and [Ca^2+^]_ER_, we hypothesized that this PM-bound protein might facilitated tethering between the ER and the PM, thereby contributing to enhanced SOCE and ER Ca^2+^ storage. To test this, we co-transfected HeLa cells with KCNT1 and GFP-MAPPER, an ER-PM contact indicator ^33^, and imaged the basal surface. We observed that the punctate distribution of KCNT1 could colocalize with the MAPPER signal (Figure 1N). This is not a universal pattern of membrane proteins considering that overexpressed E-cadherin displayed a distinct distribution pattern (Figure S1N), suggesting besides its role in K^+^ transport, KCNT1 might take part in facilitating ER-PM connection.

### K^+^ channel-mediated hyperpolarization potentiates SOCE, whereas ER Ca^2+^ modulation is dependent on the KCNT1 cytoplasmic tail

*KCNT1:R428Q* is a well-characterized epilepsy-related *GOF* mutation that results in larger outward current and increased neuronal firing rates ^34^. Using the voltage-sensitive dye DiBAC_4_(3) to measure relative resting membrane potential ^35^, we confirmed that KCNT1^R428Q^-expressing HEK293T cells were more hyperpolarized than KCNT1^WT^ cells (Figure S2A, B). Upon depolarization with 50 mM KCl, KCNT1^R428Q^ demonstrated a further increase in SOCE intensity compared to KCNT1^WT^. Intriguingly, however, it did not significantly alter the magnitude of ER Ca^2+^ release (Figure S2C, Figure 2A-C). This suggests while SOCE is sensitive to channel-induced hyperpolarization, the regulation of ER Ca^2+^ release may be independent of specific *GOF* status.

**Figure 2.**
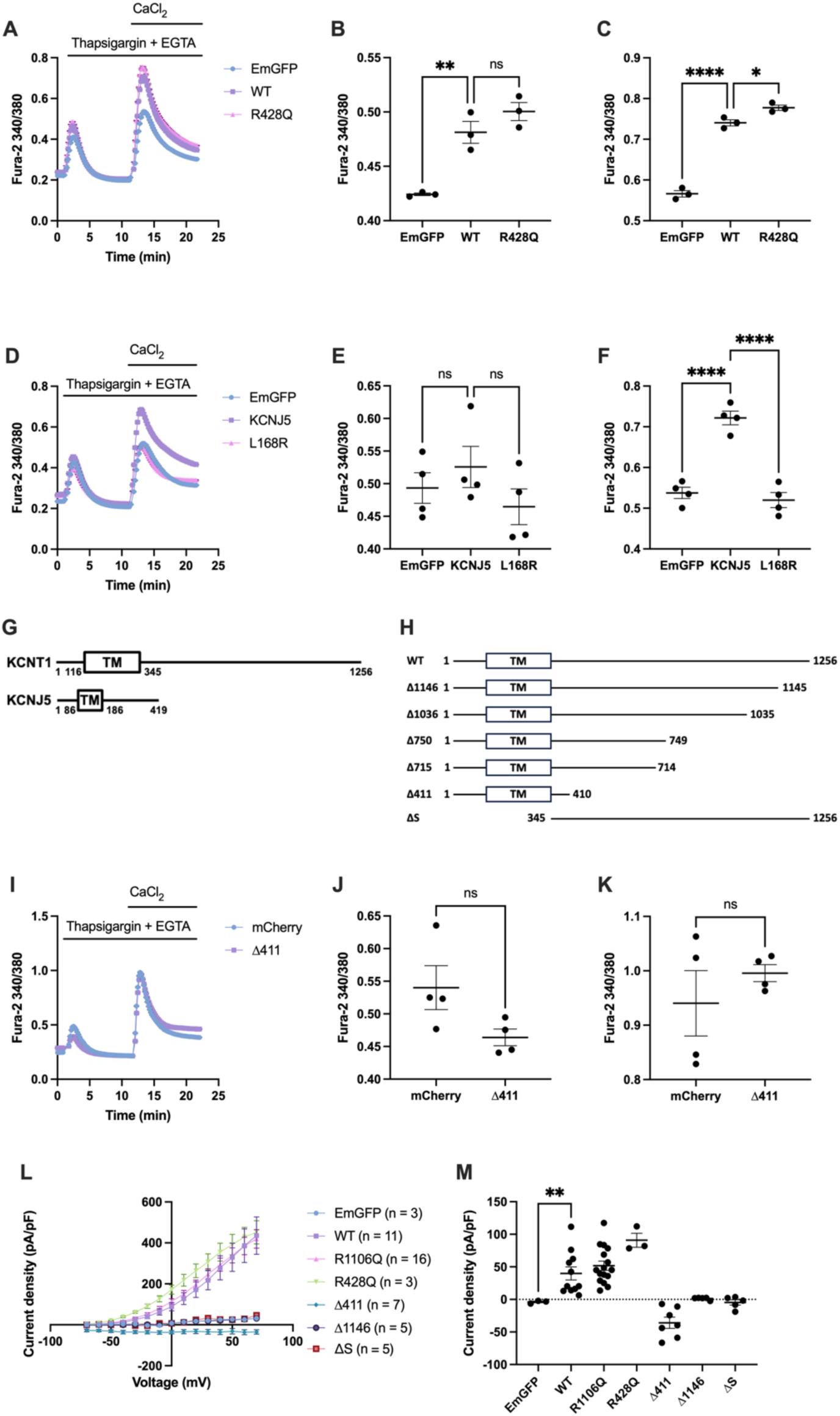
The cytoplasmic tail of KCNT1 is essential for Ca^2+^ homeostasis modulation. (A) Representative traces of Fura-2 ratio changes under 50 mM extracellular [K^+^] conditions. (B-C) Quantification of ER Ca^2+^ release (B) and SOCE intensity (C) in high [K^+^] buffer. KCNT1^R428Q^ significantly increased SOCE intensity compared to KCNT1^WT^ but did not further enhance ER Ca^2+^ release. (D) Representative Fura-2 ratio traces of HEK293T cells overexpressing KCNJ5^WT^ or the KCNJ5^L168R^ mutant. (E-F) Quantification of ER Ca^2+^ release (E) and SOCE intensity (F). While KCNJ5^WT^ significantly elevated SOCE intensity, it had no significant effect on ER Ca^2+^ release. (G) Schematic comparison of the protein structures of KCNT1 and KCNJ5, highlighting the significantly longer cytoplasmic tail of KCNT1. TM: transmembrane domains. (H) Schematic representation of KCNT1^WT^ and its various C-terminal truncated variants. (I) Representative Fura-2 ratio traces of HEK293T cells overexpressing the KCNT1^Δ411^ truncation mutant. (J-K) Quantification of ER Ca^2+^ release (J) and SOCE intensity (K). Overexpression of the cytoplasmic tail-truncated KCNT1^Δ411^ failed to alter either parameter. (L) Current-voltage (I-V) relationships of HEK293T cells overexpressing EmGFP, KCNT1^WT^, and its variants. (M) Quantification of steady-state current density recorded at −20 mV. KCNT1^WT^ significantly increased outward currents compared to the EmGFP control. \**p < 0.05*.

To further test whether altered Ca^2+^ homeostasis is driven solely by electrostatic changes, we examined KCNJ5, another K^+^ channel, and its *LOF* mutation, KCNJ5^L168R36^. Although KCNJ5 hyperpolarized the membrane potential and significantly elevated SOCE intensity, it failed to elicit a statistically significant increase in ER Ca^2+^ release (Figure S2D, Figure 2D-F). Notably while KCNJ5 localized at ER-PM junctions, its presence was insufficient to replicate KCNT1’s effect on ER Ca^2+^ release (Figure S2E).

To find out the structural basis for this difference, we compared the sequences of KCNT1 and KCNJ5. KCNT1 possesses an exceptionally large cytoplasmic tail compared to the relatively short cytoplasmic tail (Figure 2G). We hypothesized that the bulky cytoplasmic tail is the key determinant for KCNT1-mediated ER Ca^2+^ modulation. To test this, we generated a C-terminal truncated construct, KCNT1^Δ411^, which comprises of the first 410 amino acids, and lacks the most of the cytoplasmic domain (Figure 2H). Unlike the full-length protein, KCNT1^Δ411^ failed to alter either ER Ca^2+^ release or SOCE intensity (Figure 2I-K). Noteworthily, whole-cell patch-clamp recording revealed that KCNT1^Δ411^ exhibited a constitutive “leaky” conductance, a feature not observed in other truncated variants (Figure 2L, M).

The R1106Q mutation located at the tail end of KCNT1. KCNT1 ends with a PDZ binding motif (ETQL), PDZ is a scaffold protein that can bind to other proteins and affects their surface expression ^37^. Previous research indicated that K^+^ channels’ cytoplasmic tail influences their open probability and trafficking ^38,39^. To ensure that the observed alteration of KCNT1^R1106Q^ and KCNT1^Δ411^ was not due to impaired localization, we performed IF staining and confirmed that these mutants were successfully translocated to the PM (Figure S2F).

### The cytoplasmic domain is sufficient for Ca^2+^ regulation, independent of ion channel conductance

Since KCNT1^Δ411^ abolished the KCNT1-mediated Ca^2+^ flux. We further investigated the functional importance of various segments within the cytoplasmic tail by generating a series of C-terminal truncations (Figure 2H). Truncated KCNT1s with more preserved cytoplasmic sequence, such as KCNT1^Δ1146^, better retained the ability to modulate ER Ca^2+^ release, and SOCE was more sensitive to C-terminal impairment (Figure 3A-C). By pooling the z-scores from multiple truncation experiments and plotting them against the protein sequence length, we identified two potential trend curves. These curves suggest a rapid decline in SOCE intensity upon C-terminal impairment, whereas the impact on ER Ca^2+^ release followed a more moderate decline, potentially exhibiting a plateau between residues E700 and M900. (Figure 3D).

**Figure 3.**
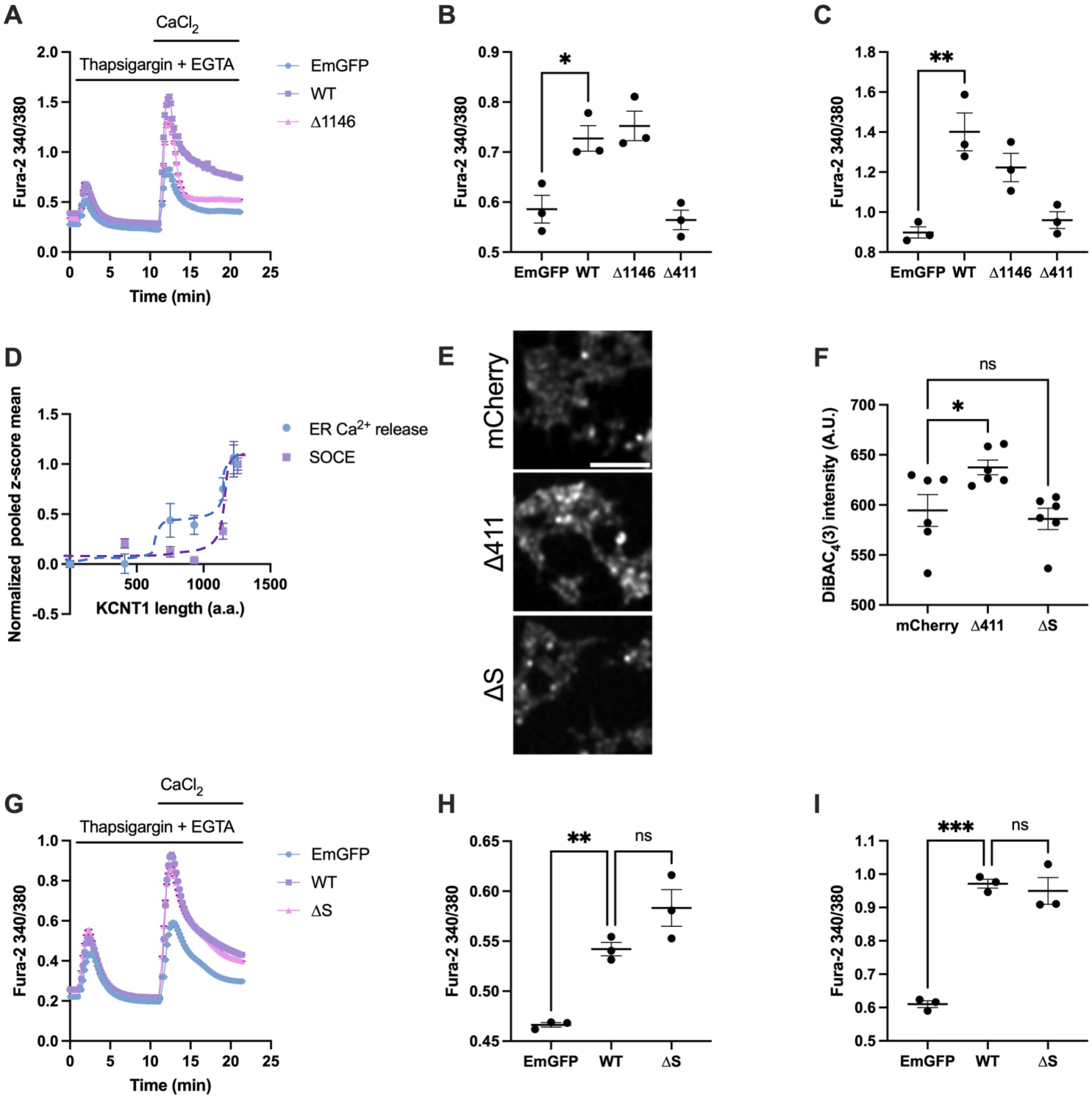
Overexpression of the KCNT1 cytoplasmic tail alone upregulate ER Ca^2+^ release and SOCE. (A) Representative Fura-2 ratio traces of HEK293T cells overexpressing the KCNT1^Δ1146^ mutant. (B-C) Quantification of ER Ca^2+^ release (B) and SOCE intensity (C). The KCNT1^Δ1146^ variant did not significantly alter ER Ca^2+^ release but exhibited a trend toward reduced SOCE compared to KCNT1^WT^. (D) Pooled average z-scores for ER Ca^2+^ release and SOCE across various KCNT1 truncation mutants. Dotted lines represent hypothetical trend curves correlating Ca^2+^ flux alterations with KCNT1 protein length. (E) Representative fluorescence images of HEK293T cells stained with DiBAC_4_(3). Cells overexpressing KCNT1^Δ411^ exhibited increased fluorescence, whereas the signal in cells overexpressing KCNT1^ΔS^ was comparable to the mCherry control. Scale bar: 100 µm. (F) Quantification of DiBAC_4_(3) fluorescence intensity, demonstrating that KCNT1^ΔS^does not significantly alter the resting membrane potential. (G) Representative Fura-2 ratio traces demonstrating that the Ca^2+^ dynamics of KCNT1^ΔS^-expressing cells closely resemble those of KCNT1^WT^. (H-I) Overexpression of KCNT1^ΔS^ alone was sufficient to significantly enhance both the peak of ER Ca^2+^ release (H) and SOCE intensity (I) comparable with KCNT1^WT^. \**p < 0.05*.

To definitively rule out the influence of K^+^ conductance, we generated a transmembrane domain-deleted construct (KCNT1^ΔS^), comprising residues Q345 to L1256, to isolate the effects of the cytoplasmic tail. (Figure 2H). Patch-clamp and DiBAC_4_(3) measurements confirmed that KCNT1^ΔS^ exhibited electrophysiological characteristics similar to the mCherry control (Figure 2L, M, Figure 3E, F). Without transmembrane domain as anchorage, KCNT1^ΔS^ dispersed evenly throughout the cell and significantly upregulated ER Ca^2+^ release and SOCE (Figure S3B, Figure 3G-I). To further explore the role of membrane localization, we attached PM-targeting signal peptides, Lck or CAAX ^40,41^, to KCNT1^ΔS^ (Figure S3A, B). The N-terminally anchored Lck-KCNT1^ΔS^ demonstrated more consistent upregulation ER Ca^2+^ release and SOCE upregulation (Figure S3C-E). These results imply that both the physical attachment to the PM and the spatial orientation of the KCNT1 cytoplasmic tail are critical determinants of its Ca^2+^ regulatory function.

### The KCNT1 cytoplasmic tail expands the lamellar ER and enhances ER-PM junction contact formation

Our data suggested that, in addition to upregulating ER Ca^2+^ release and SOCE, KCNT1 localized to ER-PM junctions. The SOCE complex comprises ER-resident STIM1 and PM-resident Orai1; therefore, we investigated whether the ER structurally participates in KCNT1-mediated Ca^2+^ modulation. Upon labeling the ER with ER-Tracker Green, we observed that KCNT1 overexpression led to a more uniform dispersion throughout HeLa cells compared to the mCherry control (Figure 4A). The ER can be classified into sheet-like (lamellar) and tubule-like structures; upon ER Ca^2+^ depletion, the regions closely apposed to the PM predominantly consist of STIM1-enriched flattened ER sheets ^42,43^. By categorizing ER-Tracker fluorescence intensity, we found that a higher proportion of bright signals correlated with lamellar ER composition (Figure S4A, Figure 4B). To validate this, we evaluated the distribution of cytoskeleton-associated protein 4 (CLIMP-63), an established lamellar ER marker ^44^. In KCNT1-overexpressing cells, CLIMP-63 covered a significantly larger relative area than in EmGFP-overexpressing cells (Figure 4C, D). Further analysis revealed that KCNT1^WT^, KCNT1^R1106Q^ and KCNT1^ΔS^ all increased the proportion of lamellar ER. Notably, KCNT1^WT^ exhibited a significantly higher proportion than the other variants, indicating that KCNT1 can influence ER distribution, a property likely associated with its cytoplasmic tail (Figure 4E).

**Figure 4.**
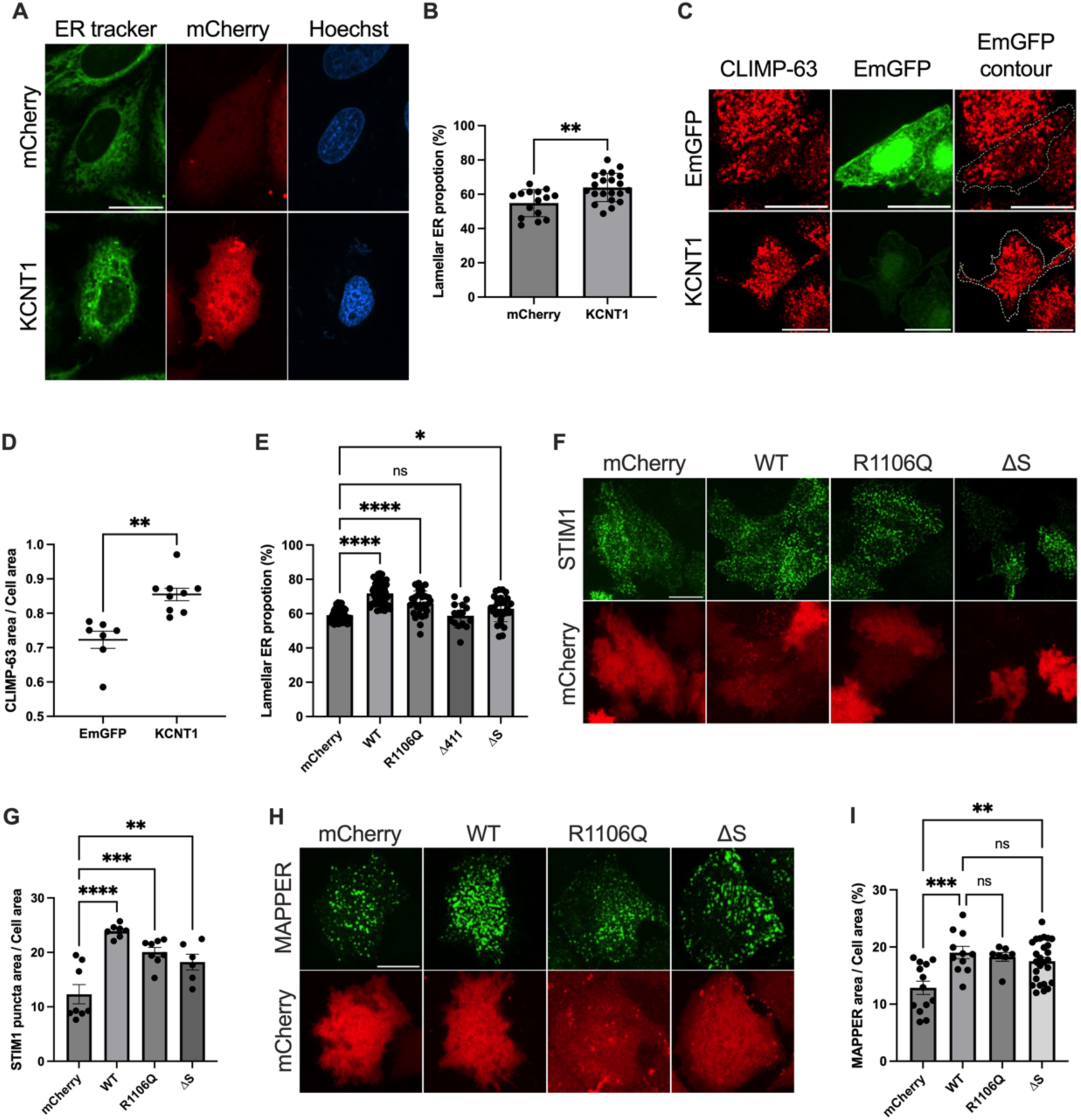
The cytoplasmic tail of KCNT1 alone is sufficient to expand the lamellar ER and facilitate ER-PM tethering. (A) Representative images of ER distribution labeled by ER-Tracker Green in HeLa cells. (B) KCNT1-overexpressing cells significantly increased the lamellar ER proportion compared to the mCherry control. (C) Representative images of CLIMP-63 signal in HeLa cells. (D) The relative CLIMP-63-positive area was significantly larger in KCNT1-overexpressing cells than in the EmGFP control. (E) Quantification of the lamellar ER proportion across different KCNT1 variants based on ER-Tracker Green signals. KCNT1^WT^, KCNT1^R1106Q^, and KCNT1^ΔS^ significantly increased the lamellar ER proportion, with KCNT1^WT^ exhibiting a significantly higher proportion than KCNT1^R1106Q^ and KCNT1^ΔS^. (F) Representative images of YFP-STIM1 distribution in HeLa cells following ER Ca^2+^ store depletion. (G) Both full-length KCNT1 variants and KCNT1^ΔS^ significantly increased relative STIM1 puncta-positive area. (H) Visualization of ER-PM contacts in HeLa cells. (I) Both full-length KCNT1 variants and KCNT1^ΔS^ significantly expanded the relative contact area. Scale bar: 20 µm. \**p < 0.05*. Image contrast in (A), (F), and (H) was adjusted independently.

To examine this structural influence further, we visualized KCNT1 and STIM1 colocalization in ER Ca^2+^ depleted HeLa cells. Unlike the evenly dispersed mCherry control, KCNT1 exhibited a more punctate distribution that overlapped with STIM1 puncta (Figure S4B). Although *KCNT1* mRNA level did not strictly correlate with those of *STIM1*, cells overexpressing KCNT1^WT^, KCNT1^R1106Q^, and KCNT1^ΔS^ all displayed a significantly higher relative STIM1 puncta area compared to the mCherry control. Among the three variants, KCNT1^WT^ displayed the highest puncta are proportion (Figure S4C, Figure 4F, G). The increase in STIM1 puncta area proportion, which is uncoupled from transcriptional upregulation, aligns with the expanded lamellar ER area, the primary site where STIM1 translocates to juxtapose with Orai1 and form functional SOCE complexes.

Given the ability of KCNT1^ΔS^ to increase STIM1 puncta, we hypothesized the KCNT1 cytoplasmic domain increases the structural accessibility of the ER to the PM. We evaluated ER-PM contact in response to different KCNT1 genotypes in HeLa cells. Both full-length KCNT1 and KCNT1^ΔS^ significantly increased the percentage of the overall ER-PM junction area compared to the mCherry control (Figure 4H, I). Collectively, these data indicate that the cytoplasmic tail of KCNT1 facilitates the tethering and approximation of the ER and the PM.

### The acidic and basic motifs within the KCNT1 cytoplasmic tail coordinate ER-PM junction formation and ER Ca^2+^ homeostasis

PM proteins can interact with ER targets, and in these interactions, charged motifs play a crucial role in binding with lipid and proteins ^45,46^. The *KCNT1:R1106Q* mutation replaces a positively charged arginine with a neutral glutamine. With this charge replacement, we not only observed a mild decline in ER Ca^2+^ release, [Ca^2+^]_ER_ and lamellar ER proportion were both significantly disrupted. Prompted by this, we analyzed the net charge distribution of KCNT1 and KCNJ5 sequences using a rolling window analysis. We identified several prominent charge-dense segments within the KCNT1 cytoplasmic tail, whereas the cytoplasmic tail of KCNJ5 lacks comparable basic clusters (Figure S5A).

To test whether these charged regions are required for Ca^2+^ modulation, we generated KCNT1^Δ715^ which lacks a major acidic cluster presents in KCNT1^Δ750^, which can still partially upregulate ER Ca^2+^ release (Figure 2H, 3D, Figure S5A). KCNT1^Δ715^ failed to increase either ER Ca^2+^ release or SOCE intensity compared to the EmGFP control (Figure 5A-C). Furthermore, co-transfection with GFP-MAPPER in HeLa cells revealed that the relative ER-PM junction area in KCNT1^Δ715^–expressing cells remained at baseline levels, suggesting that this this acidic region is essential for establishing a structural connection with the ER (Figure 5D, E).

**Figure 5.**
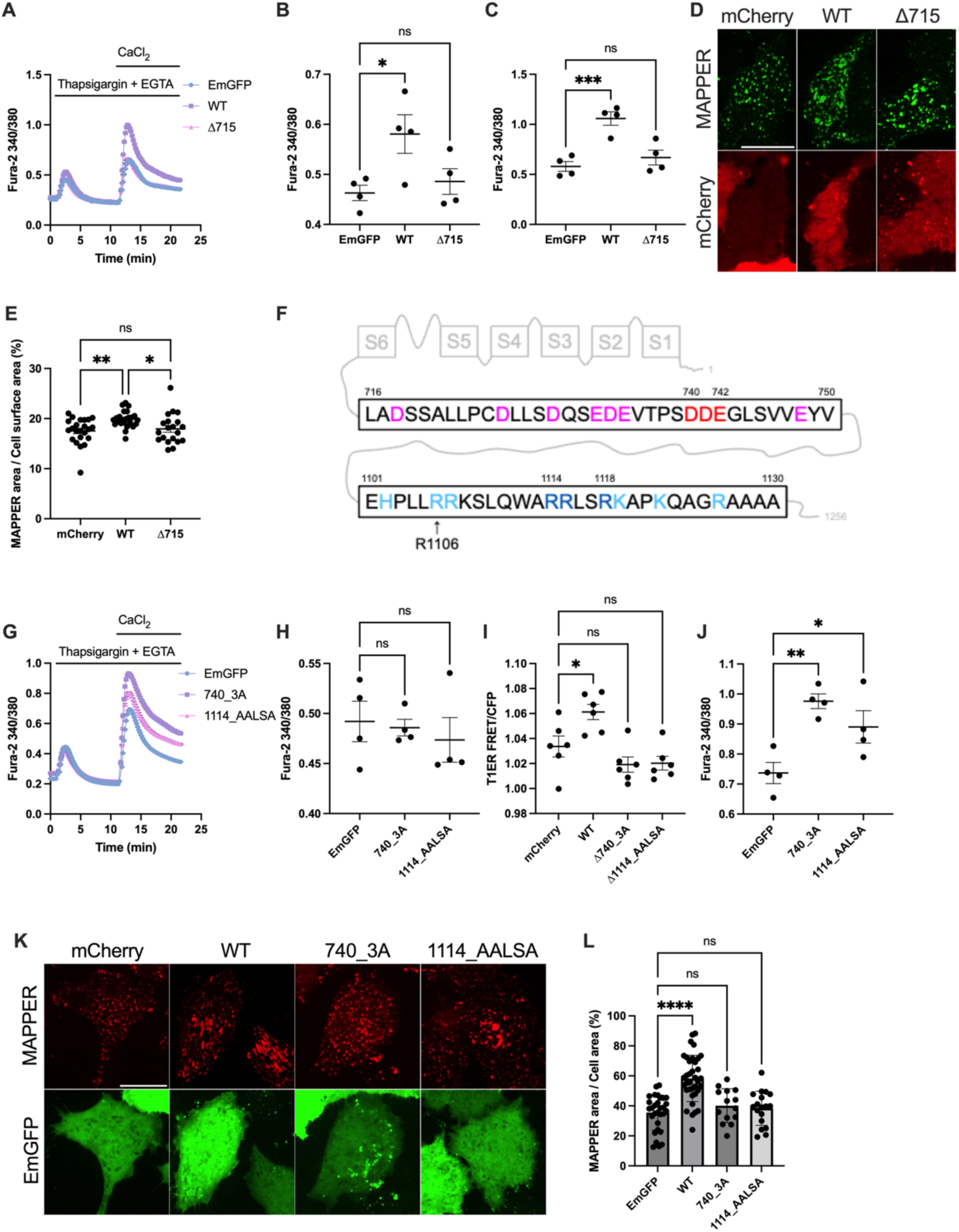
Charged motifs within the KCNT1 cytoplasmic tail are influential in ER Ca^2+^ homeostasis and ER-PM junction formation. (A) Representative Fura-2 ratio traces showed that the Ca²⁺ dynamics in KCNT1^Δ715^–overexpressing cells were comparable to the EmGFP control. (B-C) Quantification of ER Ca²⁺ release (B) and SOCE intensity (C). KCNT1^Δ715^ failed to increase either parameter. (D) Representative images of GFP-MAPPER distribution in HeLa cells overexpressing KCNT1^Δ715^. (E) Quantification showed that KCNT1^Δ715^ did not increase the ER-PM contact area proportion compared to the mCherry control. (F) Sequence analysis of the KCNT1 cytoplasmic tail. Magenta and light blue characters represent acidic and basic residues, respectively. Residues highlighted in red and blue were targets for alanine substitution. (G) Representative Fura-2 ratio traces of KCNT1^740_3A^ and KCNT1^1114_AALSA^. (H) Neither KCNT1^740_3A^ nor KCNT1^1114_AALSA^ significantly altered ER Ca²⁺ release. (I) Measurement of the T1ER FRET/CFP ratio. Both KCNT1^740_3A^ and KCNT1^1114_AALSA^ abolished the KCNT1-mediated increase in ER Ca²⁺ storage. (J) Both KCNT1^740_3A^ and KCNT11^114_AALSA^ still upregulated SOCE. (K) Distribution of mCherry-MAPPER at the basal surface of HeLa cells. (L) Neither KCNT1^740_3A^ nor KCNT1^1114_AALSA^ significantly changed the MAPPER area proportion compared to the EmGFP control. Scale bars: 20 µm. \**p < 0.05*. Image contrast was adjusted independently.

Several studies have describing ER proteins interacting with their targets through electrostatic attraction. For example, the conserved B site of Sec24 plays an important role in interacting with DxE signal sequence by forming salt bridges ^47,48^. In KCNT1, between E715 and Y749, we identified an acidic 740-DDE-742 sequence that potentially serve as a binding site for ER-resident proteins (Figure 5F, Figure S5B).

Given that KCNT1 also possesses several basic clusters, and that *KCNT1:R1106Q* mutation impaired ER Ca^2+^ homeostasis, we investigated the roles of these positively charged motifs in Ca^2+^ homeostasis. Clusters of basic residues often facilitate interactions with phosphatidylinositol 4,5-bisphosphate (PIP_2_), a lipid involved in various ion channel activities ^49,50^. In KCNT1, we identified a 1114-RRLSR-1118 sequence located near the C-terminus and proximal R1106 (Figure 5F, Figure S5C). Based on these observations, we performed site-directed mutagenesis and generated two mutations: KCNT1^740_3A^ (D740A/D741A/E742A) and KCNT1^1114_AALSA^ (R1114A/R1115A/R1118A). Unlike KCNT1^WT^, neither KCNT1^740_3A^ nor KCNT1^1114_AALSA^ increased ER Ca^2+^ release or the [Ca^2+^]_ER_ (Figure 5G, I). However, these substitutions did not abolish KCNT1-mediated SOCE potentiation (Figure 5G, J).

Although we previously demonstrated that KCNT1^WT^ colocalize with the MAPPER signal, examination of the mutant distributions revealed that most punctate KCNT1^740_3A^ failed to overlay with MAPPER. In contrast KCNT1^1114_AALSA^ mutation appeared less impacted in this regard (Figure S5D). To examine how KCNT1^740_3A^ and KCNT1^1114_AALSA^ influence ER-PM junctions, we co-transfected HeLa cells with mCherry-MAPPER and the respective KCNT1 variants. Quantification of concentrated MAPPER area ratio showed that both KCNT1^740_3A^ and KCNT1^1114_AALSA^ failed to increase relative MAPPER signal area compared to the EmGFP control (Figure 5K, L). Together, these results indicate that charge-carrying motifs on the cytoplasmic tail of KCNT1 are crucial for facilitating ER Ca^2+^ homeostasis and the formation of ER-PM junctions.

### The KCNT1 cytoplasmic tail shortens the FPD and increases the spontaneous beating rate of hiPSC-CMs

Having demonstrated that KCNT1, specifically through its cytoplasmic tail, was capable of changing cellular Ca^2+^ homeostasis, more specifically, ER Ca^2+^ release and SOCE in non-excitable cell, we next sought to evaluate whether these effects transfer to excitable cells. Given that the *KCNT1:R1106Q* mutation was originally identified in a BrS patient, we investigated KCNT1’s effect on the electrophysiology of hiPSC-CMs.

Cells were seeded on MEA chips at high density. To evaluate changes in the FPD, waveforms were recorded under paced conditions (125 ms stimulation at 300 ms interval; Figure 6A). While the mCherry control and KCNT1^Δ411^ groups exhibited a prolonged FPD compared to their pre-transduction baseline, KCNT1^WT^, KCNT1^R1106Q^, and KCNT1^ΔS^ all significantly shortened the FPD (Figure S6A, Figure 6B).

**Figure 6.**
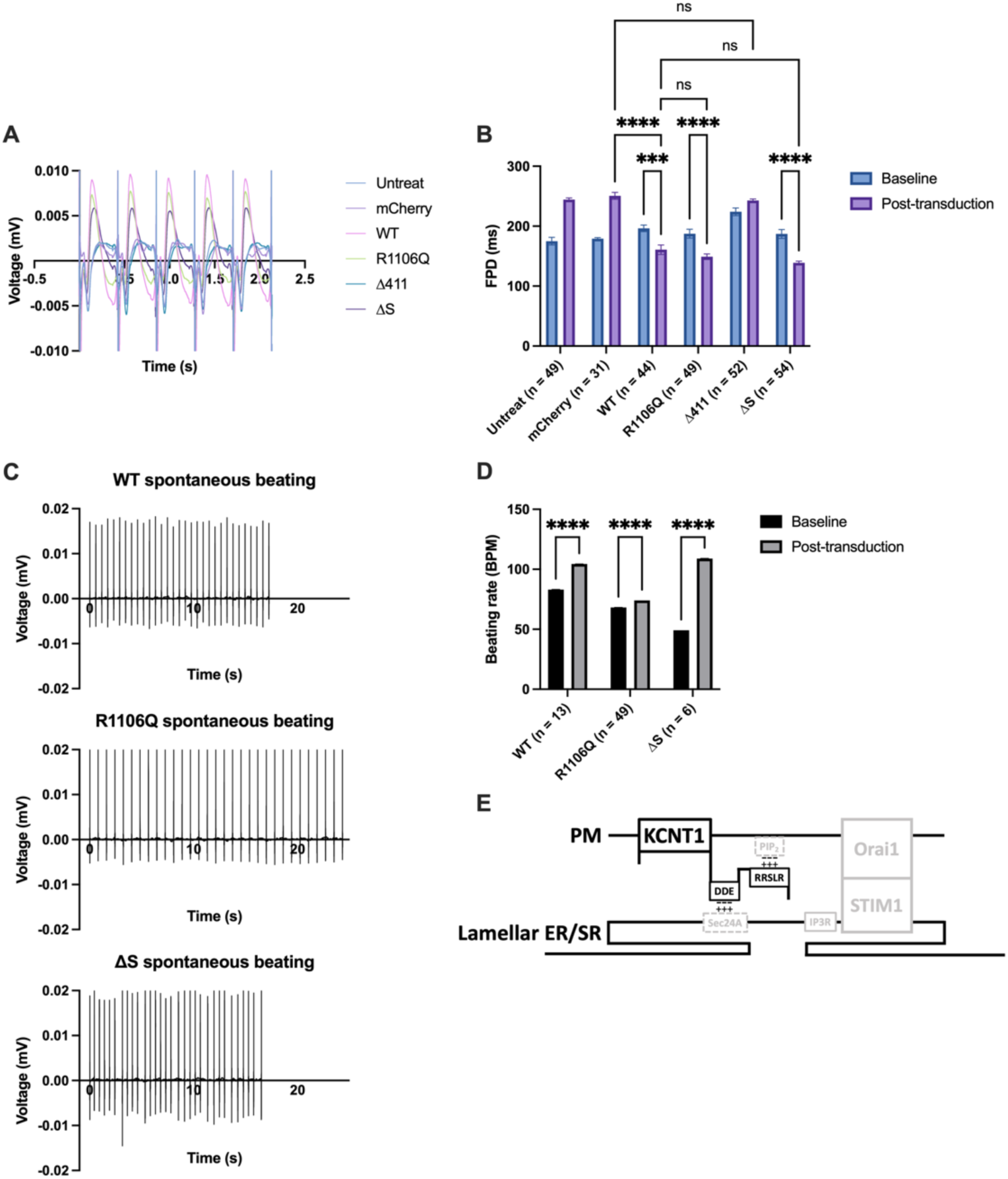
The KCNT1 cytoplasmic tail is sufficient to modulate cardiomyocyte electrophysiology by shortening FPD and accelerating spontaneous beating rates. (A) Representative field potential waveforms of hiPSC-CMs transduced with KCNT1^WT^ or its variants under paced conditions. (B) Quantification of the FPD before and after lentiviral transduction. Overexpression of KCNT1^WT^, KCNT1^R1106Q^, and KCNT1^ΔS^ significantly shortened the FPD, which was lengthened by the mCherry control and KCNT1^Δ411^. (C) Representative voltage traces showed the spontaneous beating activity of transduced hiPSC-CMs. (D) Comparison of spontaneous beating rates before and after transduction. KCNT1^WT^, KCNT1^R1106Q^, and KCNT1^ΔS^ all significantly accelerated the beating rate. (E) Proposed model for the non-conducting role of KCNT1 in intracellular Ca²⁺ regulation. The charged motifs distributed along the KCNT1 cytoplasmic tail serve as structural anchors that bridge the ER and the PM, facilitating ER-PM junction formation and enhancing ER Ca²⁺ release. n: the number of electrodes. \*\*\**p < 0.001*.

In addition to FPD measurements, we accessed spontaneous beating activity (Figure 6C). Overexpression of KCNT1^WT^, KCNT1^R1106Q^, and KCNT1^ΔS^ consistently increased the spontaneous beats rate (beats per minute, BPM; Figure 6D). Notably, the KCNT1^ΔS^ group exhibited a significantly higher fold change in beating rate compared to the other groups, although this may be attributed to a lower baseline beating rate prior lentiviral transduction (Figure S6B). Conversely, the magnitude of the beating rate increase in the KCNT1^R1106Q^-overexpressing cells was significantly attenuated compared to the other two groups.

Collectively, these results demonstrate that KCNT1 and its variants can modulate cardiomyocyte physiology, a function that can be recapitulated by the cytoplasmic tail alone.

## Discussion

Traditionally, it has been proposed that Kv channels influence Ca^2+^ homeostasis primarily through the hyperpolarizing effects of their rectifying K^+^ current. However, our findings regarding KCNT1’s impact on ER Ca^2+^ storage and release suggest a more complex regulatory role. Although these alteration in Ca^2+^ signaling might be interpreted as a secondary consequence of excessive Ca^2+^ influx, the colocalization of KCNT1 with ER-PM junction marker points toward a direct structural involvement. Our data support a model in which the oppositely charged motifs distributed along the KCNT1 cytoplasmic tail serve as structural anchors. In this non-conducting capacity, the cytoplasmic tail bridges the ER and the PM, facilitating the spatial organization requisite for efficient Ca^2+^ handling (Figure 6E). This interpretation is further substantiated by our experiments utilizing KCNT1^ΔS^, which lacks the transmembrane domains but retains the ability to modulate Ca^2+^ homeostasis. The observation that PM-anchoring via Lck further enhances this effect underscores the importance of the cytoplasmic tail’s physical proximity to the membrane. This study initially sought to elucidate the pathophysiological alterations induced by the BrS-associated *KCNT1:R1106Q* mutation. To evaluate the structural impact of this point mutation, *in silico* analysis using DynaMut2 was performed ^51^. This analysis predicted a stability change (ΔΔG^Stability^) of −0.52 kcal/mol, indicating to a destabilized state, alongside the potential formation of a new polar bond between Q1106 and K1108.

Functionally, the *KCNT1:R1106Q* mutation results in a moderate decline in ER Ca^2+^ release, a phenomenon supported by a significantly downregulated T1ER FRET ratio. Conversely, KCNT1^R1106Q^ further upregulates SOCE intensity, which aligns with the paradigm that BrS-associated K^+^ channelopathies are typically *GOF* mutations. Surprisingly, however, KCNT1^R1106Q^ does not significantly amplify the outward current. While a marginal increase was observed, its magnitude was negligible when compared to the pronounced current amplification seen in the epilepsy-related *KCNT1:R428Q* mutant.

Although the absence of substantial increase in outward current may appear counterintuitive for a *GOF* variant, this subtlety is physiologically plausible. Considering that the average age of the first arrhythmic event in BrS patients ranges from 39.5 to 49 years ^52^, suggesting *KCNT1:R1106Q* may not provoke drastic, immediate phenotypic changes.

An intriguing finding of this study is profound impact of KCNT1 on ER Ca^2+^ homeostasis. While ER-PM junctions and Ca^2+^ signaling are known to mutually influence each other ^53^, the KCNT1-mediated enhancement of ER Ca^2+^ release cannot be solely attributed to colocalization at ER-PM contact, as the overexpression of KCNJ5 failed to produce a comparable increase. We observed that progressive truncation of the cytoplasmic tail disrupted Ca^2+^ modulation. These data suggest that specific motifs within the cytoplasmic tail directly orchestrate Ca^2+^ flow, independent of the channel’s electrostatic status.

Polyacidic sequences are implicated in binding to ER-resident proteins. Our experiments utilizing KCNT1^Δ715^ highlighted the necessity of negatively charged motifs for KCNT1 function. A parallel can be drawn with Kv2.1, where phosphorylated Kv2.1 interacts with VAMP-associated protein (VAP) via its FFAT motif, facilitating the coupling of LTCCs to ryanodine receptor (RyR) to mediate LTCC opening and Ca^2+^ spark generation in neurons ^54^. Although KCNT1 lacks a canonical FFAT motif sequence, the interaction between FFAT motif and VAP is fundamentally initiated by non-specific electrostatic binding ^55^. This implies that cytoplasmic proteins harboring polyacidic sequences may still be capable of interacting with VAP or similar ER tethering proteins. Consistent with this hypothesis, KCNT1^740_3A^ demonstrated that disrupting the polyacidic 740-DDE-742 sequence significantly impaired both ER Ca^2+^ release and ER-PM contact.

Conversely, the role of positively charged motifs in mediating protein-PM interactions should also be considered, particularly that R1106 is flanked by multiple basic amino acids, and the *KCNT1:R1106Q* mutation downregulates [Ca^2+^]_ER_ while exhibiting a tendency to reduce ER Ca^2+^ release. For instance, the polybasic motif within the STIM1 cytoplasmic tail binds to negatively-charged phosphoinositide lipids in the PM ^53^. Our experiments on KCNT1^1114_AALSA^ demonstrate the essential role of the polybasic sequence at the distal end of the KCNT1 cytoplasmic tail in maintaining these structural connections.

Our MEA assay revealed that both KCNT1^WT^, KCNT1^R1106Q^ shorten the FPD in hiPSC-CMs. The reduction is anticipated, as overexpressed KCNT1 amplified outward current compared to the EmGFP control, thereby accelerating the repolarization process. Previous studies have established that elevated extracellular and intracellular [Ca^2+^] enhances plateau height of APs and promotes Ca^2+^-induced Ca^2+^ release (CICR) from the SR, which subsequently facilitate repolarization and decreases AP duration ^56,57^. Notably, KCNT1^ΔS^ also modulates cardiomyocyte AP. The FPD shortening induced by KCNT1^ΔS^ closely aligns with its SR Ca^2+^ release dynamics, providing novel insights into how the KCNT1 cytoplasmic tail modulates cardiac rhythm.

Consistent with our findings that KCNT1 upregulates [Ca^2+^]_ER_, SR Ca^2+^ overload is a well-known trigger for spontaneous SR Ca^2+^ release. This aberrant generates depolarizing membrane currents, in which Na^+^-Ca^2+^ exchangers (NCX) is implicated, give rise to delayed afterdepolarizations (DADs), and eventually initiates tachyarrhythmias ^58,59^.

Intriguingly, although KCNT1^R1106Q^-expressing hiPSC-CMs exhibited a shortened FPD comparable to the KCNT1^WT^ and KCNT1^ΔS^ groups, their post-transduction acceleration in beating rate was attenuated. This attenuation may be linked to the observation that the KCNT1:R1106Q mutation results in a diminished upregulation of [Ca^2+^]_ER_ and a marginal decrease in ER Ca^2+^ release compared to the wild-type. Our findings regarding KCNT1^1114_AALSA^ offer a plausible explanation, given that this polybasic substitution occurrs immediately downstream of R1106. The loss of ER Ca^2+^ modulation in KCNT1^1114_AALSA^, a function that KCNT1^R1106Q^ partially retains, may be attributed to the differing number of basic amino acids substituted. Consequently, this partial *LOF* phenotype concerning SR Ca^2+^ handling may underlie the attenuated beating rate acceleration. Furthermore, deadly ventricular arrhythmias in BrS are frequently initiated by nocturnal bradycardia ^60^, we hypothesize that the arrhythmogenic mechanism driven by the *KCNT1:R1106Q* mutation is not primarily dictated by subtle alterations in K^+^ current. Rather, it is driven by imbalanced Ca^2+^ handling, specifically regarding SR Ca^2+^ release and storage, stemming from structural changes in the KCNT1 cytoplasmic tail. While quinidine is utilized to treat BrS and KCNT1-associated epilepsy, its lack of potency remains a clinical challenge ^61,62^. Our findings suggest that targeting the distinct charge distributions and spatial conformations of the KCNT1 cytoplasmic tail may offer an effective and mutant-specific therapeutic strategy development. Although KCNT1 was the primary focus of this investigation, the structural features underlying its modulation of Ca^2+^ homeostasis and hiPSC-CMs beating, namely the length and charge distribution of its cytoplasmic tail, are not entirely exclusive to KCNT1. This implies a broader applicability for our proposed mechanism. For instance, the human Ether-à-go-go-Related Gene (hERG) encodes a K^+^ channel frequently associated with long QT syndrome ^63^. Although its cytoplasmic tail is significantly shorter than that of KCNT1, it possesses prominent positively and negatively charged motifs. These structural elements may independently influence Ca^2+^ flow and cardiomyocyte physiology in a manner analogous to KCNT1.

## Limitation

A primary limitation of this study is that the MEA assays were conducted using hiPSC-CMs cultured in a 2D monolayer. These cultures typically lack a mature, well-developed T-tubule system running closely to the SR. Therefore, utilizing advanced methodologies, such as 3D tissue cultures that promote cell maturation and T-tubule development ^64^, will be beneficial for future investigation. Additionally, generating *KCNT1:R1106Q* knock-in animal model will be indispensable to definitively ascertain whether this specific variant is sufficient to trigger BrS *in vivo*, and to determine if targeted Ca^2+^ handling can mitigate this arrhythmogenic phenotype.

## Supporting information

Supplemental figures and methods, for the link to the file on bioRxiv

## Methods

### Detailed Methods

Detailed experimental procedures and material descriptions are provided in the Supplemental Methods. The data that support the findings of this study are available from the corresponding author upon reasonable request.

### Statistical Analysis

Data are presented as mean ± standard error of the mean. Statistical analyses were performed using GraphPad Prism software (version 11.0.0). For comparisons among three or more groups, ordinary one-way analysis of variance (ANOVA) was utilized, followed by either Tukey’s Honestly Significant Difference (HSD) or Dunnett’s post hoc tests. Brown-Forsythe and Welch ANOVA tests were employed for datasets exhibiting unequal variances or different sample sizes (Figure 2M, Figure S2D, and Figure S6B) regarding their sample size difference or data distribution. Two-way ANOVA followed by Tukey’s HSD was conducted in Figure 6. For comparisons between two independent groups, Welch’s t-test was applied. Statistical significance was defined as \**p < 0.05*, \*\**p < 0.01*, \*\*\**p < 0.001*, and \*\*\*\**p < 0.0001*.

## Acknowledgments

The authors would like to thank Mao-Ting-Hung for his technical support and for providing troubleshooting training in electrophysiological experiments. We thank the RNA Technology Platform and Gene Manipulation Core Facility (RNAi core) of the National Core Facility for Biopharmaceuticals at Academia Sinica in Taiwan for providing shRNA packaging related services.

