## Supplemental figures and methods, for the link to the file on bioRxiv for "A Brugada-related KCNT1 mutation unveils its conductance-independent activation of store-operated Ca^2+^ entry"

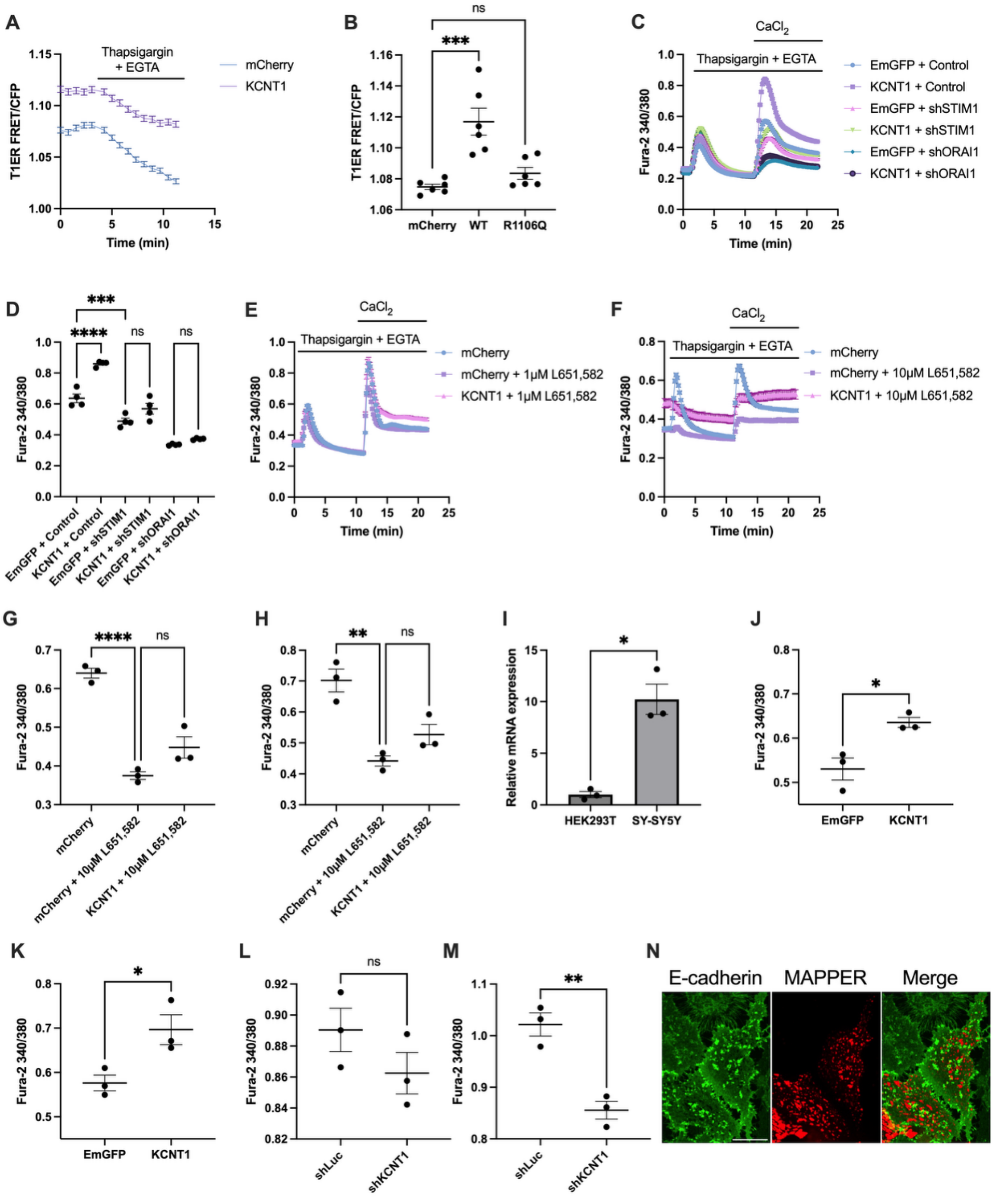


**Figure S1. Validation of $Ca^{2+}$ indicators, pharmacological inhibitors, and endogenous KCNT1 mRNA expression across cell lines.**

(A) Representative traces showing the decrease in T1ER FRET/CFP ratio following the application of thapsigargin and EGTA in HEK293T cells.

(B) Quantification of the baseline T1ER FRET/CFP ratio, demonstrated that KCNT1-overexpressing cells exhibited significantly higher ER $Ca^{2+}$ storage than $KCNT1^{R1106Q}$ and the mCherry control.

(C) Representative Fura-2 ratio traces in HEK293T cells following the knockdown of SOCE components.

(D) Knockdown of either STIM1 or Orai1 significantly reduced $CaCl_2$-induced $Ca^{2+}$ influx and abolished the difference between the EmGFP and KCNT1 groups.

(E) Representative Fura-2 ratio traces showing that 1 µM L-651,582 reduced overall $Ca^{2+}$ flow.

(F) Treatment with 10 µM L-651,582 severely disrupted $Ca^{2+}$ homeostasis.

(G-H) Quantification of ER $Ca^{2+}$ release (G) and SOCE intensity (H) in the presence of 10 µM L-651,582. Despite the baseline disruption, the relative trends in KCNT1-overexpressing cells remained observable.

(I) qPCR analysis showing significantly higher endogenous KCNT1 mRNA levels in SH-SY5Y cells compared to HEK293T cells.

(J-K) Overexpression of KCNT1 in SH-SY5Y cells significantly enhanced both ER $Ca^{2+}$ release (J) and SOCE intensity (K).

(L-M) KCNT1 knockdown did not downregulate ER $Ca^{2+}$ release significantly (L) but significantly decreased SOCE intensity (M).

(N) Representative images showing that overexpressed GFP-E-cadherin did not colocalize with mCherry-MAPPER.

Scale bar: 20 µm.

*$p < 0.05$.

Image contrast was adjusted independently.

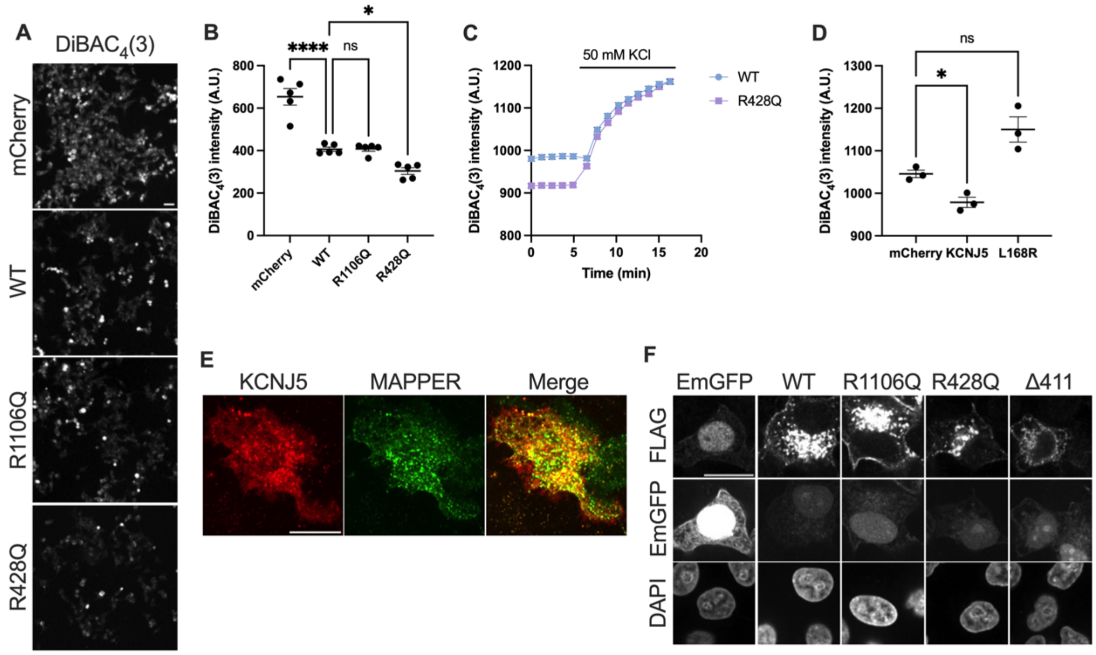


**Figure S2. Validation of membrane potential changes and subcellular localization of KCNJ5 and KCNT1 variants.**

(A) Representative images of HEK293T cells stained with DiBAC$_4$(3). Scale bar: 40 µm.

(B) Overexpression of KCNT1$^{WT}$ reduced the DiBAC$_4$(3) fluorescence compared to the mCherry control, and the KCNT1$^{R428Q}$ mutation further decreased this signal.

(C) The DiBAC$_4$(3) signal increased following the addition of 50 mM KCl, indicating cellular depolarization.

(D) KCNJ5$^{WT}$ significantly reduced the DiBAC$_4$(3) signal compared to both KCNJ5$^{L168R}$ and the mCherry control.

(E) Representative images showing that KCNJ5 colocalized with GFP-MAPPER at the basal surface of HeLa cells. Scale bar: 20 µm.

(F) FLAG-tagged KCNT1 variants were successfully translocated to the PM. Scale bar: 20 µm.

* *$p < 0.05$*.

Image contrast in (E) and (F) was adjusted independently

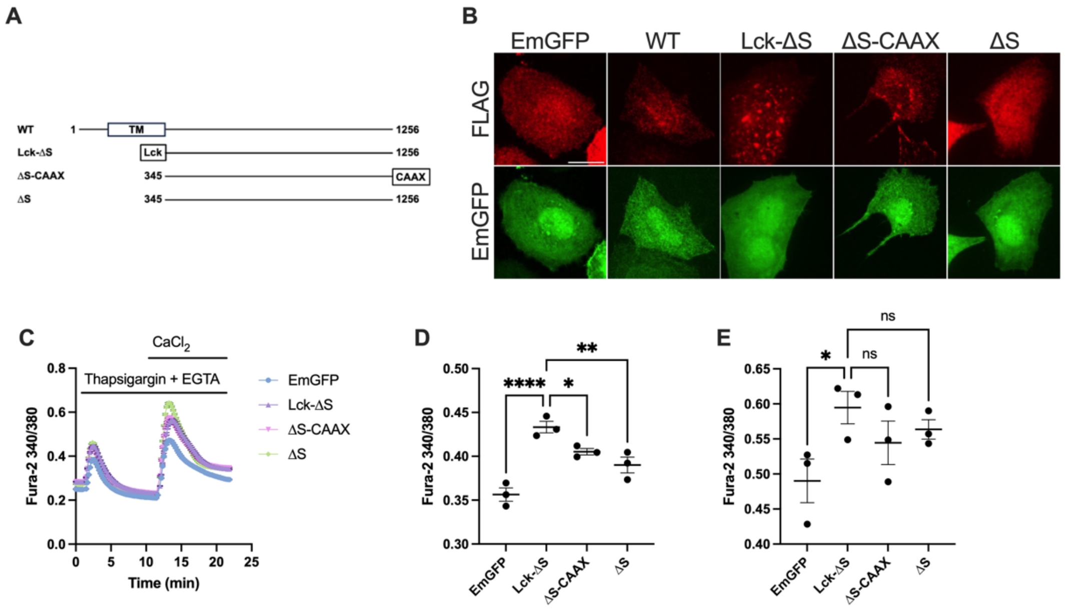


**Figure S3. PM targeting of the KCNT1 cytoplasmic tail enhances ER Ca$^{2+}$ release.**
(A) Schematic representation of the KCNT1$^{\Delta S}$ constructs fused with PM-targeting signal peptides, Lck or CAAX.
(B) Representative IF images showing the subcellular distribution of FLAG-tagged KCNT1 variants. Lck-KCNT1$^{\Delta S}$ and KCNT$^{\Delta S}$-CAAX both exhibited a more punctate distribution compared to KCNT1$^{\Delta S}$. Scale bar: 20 µm.
(C) Representative Fura-2 ratio traces of cells expressing KCNT1$^{\Delta S}$ fused with different signal peptides.
(D) The Lck-tagged KCNT1$^{\Delta S}$ significantly enhanced ER Ca$^{2+}$ release compared to the untagged KCNT1$^{\Delta S}$.
(E) Quantification of SOCE intensity. The addition of PM-targeting signal peptides did not further alter SOCE modulation.
*$p < 0.05$.
Image contrast was adjusted independently.

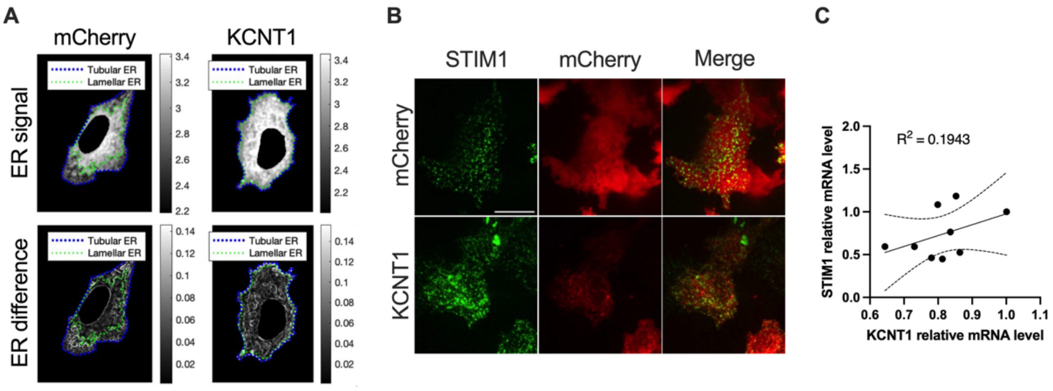


**Figure S4. Morphological classification of the ER, KCNT1-STIM1 colocalization, and transcriptional correlation.**

(A) Representative processed images illustrating the classification of ER subdomains (lamellar vs. tubular) in mCherry control (left) and KCNT1-overexpressing (right) HeLa cells. The panels display the original ER-Tracker signal, the calculated edge gradient magnitude (ER difference), and the resulting morphological classification masks (dotted lines).

(B)Representative images demonstrating the colocalization of KCNT1 with STIM1 puncta in HeLa cells following ER $Ca^{2+}$ store depletion. Scale bar: 20 µm.

(C) Correlation analysis between *KCNT1* and *STIM1* mRNA levels in SH-SY5Y cells. The data revealed a weak transcriptional correlation ($R^2$ = 0.1943).

Image contrast was adjusted independently.

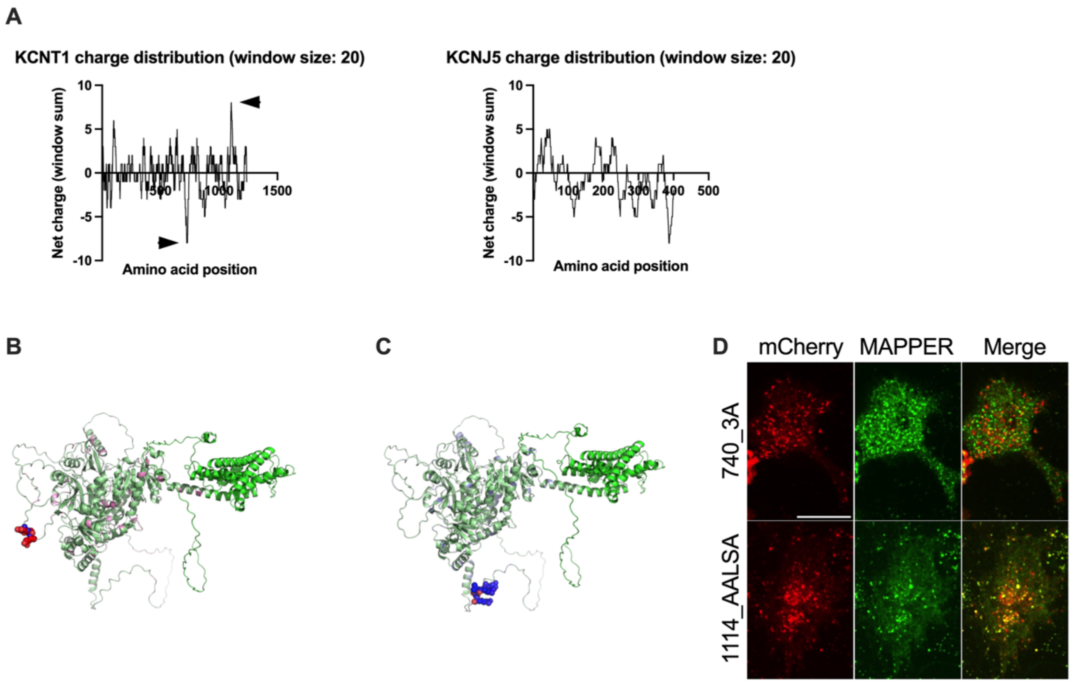


**Figure S5. Charge distribution analysis of KCNT1 and the subcellular localization of its charge-motif mutants.**

(A) Rolling window analysis of the net charge across the amino acid sequences of KCNT1 and KCNJ5. Arrowheads highlight the prominent charge-concentrated regions within the KCNT1 cytoplasmic tail.

(B-C) Structural mapping of KCNT1 depicting the distribution of acidic (B) and basic (C) amino acids. In (B), the specific 740-DDE-742 sequence was highlighted in red. In (C), the 1114-RRLSR-1118 sequence was highlighted in blue.

(D) Representative images showing the distribution of KCNT1$^{740_3A}$ and KCNT1$^{1114_AALS}$A at the basal surface of HeLa cells. KCNT1$^{740_3A}$ failed to colocalize with the ER-PM junction marker GFP-MAPPER, whereas KCNT1$^{1114_AALSA}$ retained its colocalization. Scale bar: 20 µm. Image contrast was adjusted independently.

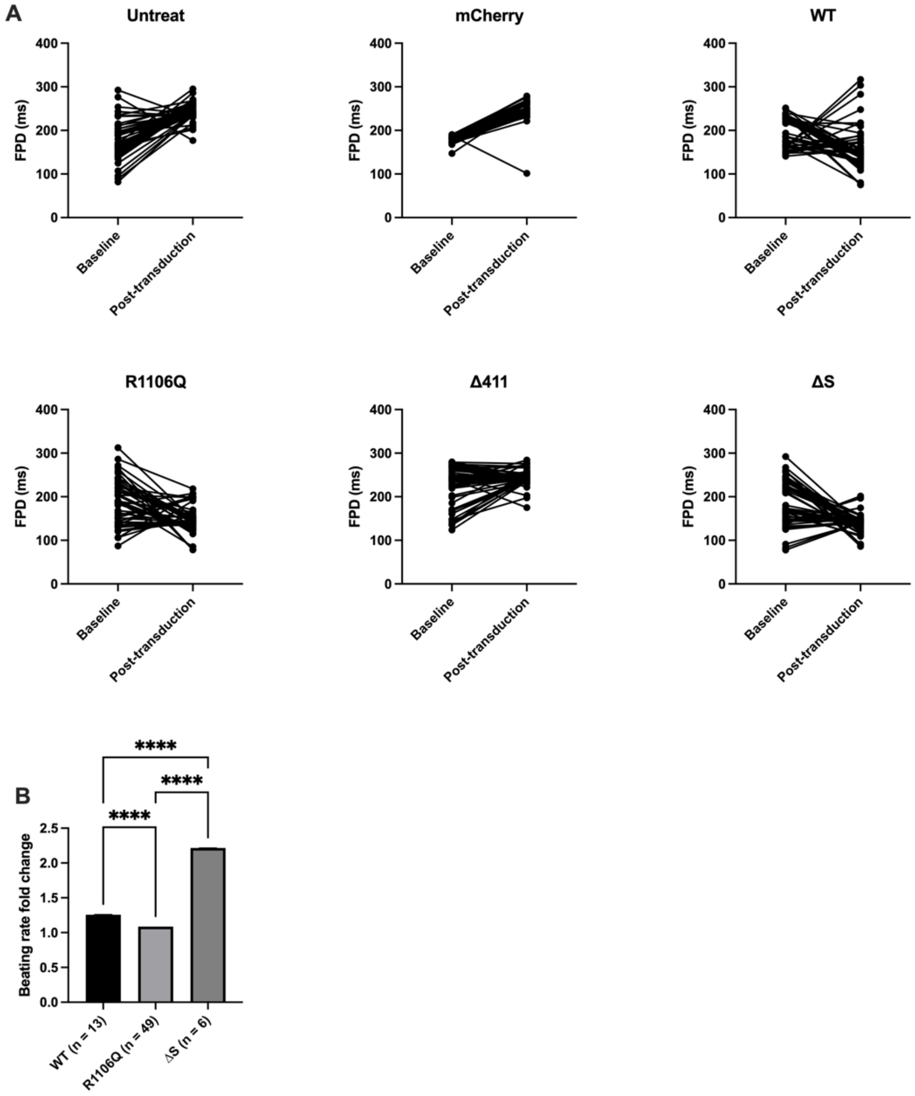


**Figure S6. Paired analysis of FPD and spontaneous beating rate fold change in hiPSC-CMs**

(A) Paired comparison of FPD recorded from individual electrodes before and two days after transduction with the indicated lentiviral variants.

(B) Fold change in the spontaneous beating rate of hiPSC-CMs relative to baseline following lentiviral transduction. n: the number of electrodes. **** *p < 0.0001*.

### Supplemental Methods

#### Cell culture

All cells were maintained in a humidified incubator at 37°C with 5% $CO_2$. HEK293T and HeLa cells were cultured in Dulbecco's Modified Eagle Medium (DMEM)/high glucose (Cytiva, SH30003) supplemented with 10% fetal bovine serum (FBS; HyClone or Biological Industries) and 1% penicillin-streptomycin (P/S; Gibco, 15140122). SH-SY5Y cells were maintained in DMEM/F12 (Gibco, 11330) with 10% FBS and 1% P/S. For cell dissociation, 0.05% Trypsin-EDTA (Gibco) in phosphate-buffered saline (PBS) was utilized.

#### $Ca^{2+}$ measurement

Cells were seeded into 96-well culture plates (Costar, 3599) at a density of 10,000–15,000 cells per well. The plates were pre-coated with collagen (60 µg/mL for HEK293T cells; 30 µg/mL for HeLa cells). The extracellular buffer (ECB) consisted of 125 mM NaCl, 5 mM KCl (Sigma, P5405), 1.5 mM $MgCl_2$ (Sigma, M4880), 10 mM glucose (Sigma, G7021), 1.5 mM $CaCl_2$ (Sigma, C5670), and 20 mM HEPES (Gibco,15630080), and pH was adjusted to 7.4. On the day of experiment, cells were loaded with 2 µM Fura-2/AM (Invitrogen, F1221), 0.02% Pluronic™ F-127 (Invitrogen, P3000MP), and 4 µM probenecid (Invitrogen, P36400) for 30 minutes at room temperature. After rinsing with ECB, cells were further incubated in ECB containing 4 µM probenecid at 37°C for 15 minutes. Ratiometric imaging was performed using a Nikon microscope with NIS-Elements software. Fura-2 fluorescence was excited at 340 and 380 nm. Following a 1-minute baseline recording, 2 µM thapsigargin (Invitrogen, T7459) and 1.5 mM EGTA (Sigma) were applied to quantify ER $Ca^{2+}$ release. After 10 minutes, 3 mM $CaCl_2$ was added to measure SOCE intensity. Data were analyzed using MATLAB.

#### Transfection

For HEK293T cells in 96-well plates, Lipofectamine™ 3000 (Invitrogen) was used according to the manufacturer's protocol: 100 ng of plasmid was mixed with 0.2 µL of P3000™ reagent and 0.3 µL of Lipofectamine™ 3000. The mixtures were incubated at room temperature for 15 minutes before application (10 µL/well). For HeLa cells, TransIT-X2 (Mirus) was employed; 100 ng of DNA was mixed with 0.3 µL of TransIT-X2 in 9 µL of Opti-MEM™ and incubated for 15 minutes prior to transfection. Cells were then incubated overnight at 37°C with 5% $CO_2$.

#### Lentivirus packaging

TransIT-LT1 (Mirus) was used for lentiviral packaging. The packaging plasmids (pCMVDR8.91 and pMD.G) and the lentiviral transfer vector were mixed in 1.5 mL Opti-MEM™ at a 9:1:10 ratio, yielding a total of 15 µg of DNA. After a 15-minute incubation at room temperature, the mixture was added to HEK293T cells seeded in a 10-cm culture dish. The following day, the medium was replaced with complete DMEM containing 1% bovine serum albumin (BSA; BioShop, 9048-46-8). Viral supernatant was collected 24 hours later and stored at 4°C, and the dish was replenished with fresh 1% BSA DMEM for a second collection the next day. The pooled supernatant was filtered through a 0.45 µm PES filter (Merk, SLHPR33RB) and mixed with Lenti-X™ Concentrator (Takara) at one-third of the supernatant volume. The mixture was incubated at 4°C for at least 30 minutes, followed by centrifugation at 1,500 × *g* for 45 minutes at 4°C. The viral pellet was resuspended in 1 mL of 1% BSA DMEM, divided into aliquots, and stored at -80°C.

**Lentivirus transduction and shRNA knockdown**

Cells were transduced with the desired lentivirus supplemented with 8 µg/mL polybrene (Santa Cruz, sc-134220). For *KCNT1* knockdown experiments, lentiviruses expressing short hairpin RNAs (shRNAs) were utilized. The specific target sequences directed against human *KCNT1* were 5'-CCTCAGCTACAAAGGCAACAT-3' and 5'-GCAAATTTAACAGCCAGTGAT-3'; *STIM1*, 5'-CGATGAGATCAACCTTGCTAA-3'; *ORAI1*, 5'-GCAACGTGCACAATCTCAACT-3'; luciferase (control), 5'-GCGGTTGCCAAGAGGTTCCAT-3'. 24 hours post-transduction, the medium was replaced with complete growth medium containing puromycin for positive selection. The puromycin concentrations used were 1 µg/mL for SH-SY5Y and 2 µg/mL for HEK293T.

**Patch-clamp recordings**

HEK293T cells were transfected overnight prior to whole-cell patch-clamp recordings. Transfected cells were re-plated onto poly-D-lysine-coated 12-mm coverslips and incubated in complete growth medium for 3 hours. Coverslips were transferred to the recording bath solution 3 minutes before data acquisition. Recordings were performed at room temperature using a HEKA EPC 10 amplifier controlled by Patchmaster software (HEKA). Data were exported as ASC files for plotting and statistical analysis in Python.

For outward $K^+$ current measurements, the bath solution contained 140 mM NaCl, 1 mM $CaCl_2$, 3 mM KCl, 29 mM glucose, and 25 mM HEPES (pH adjusted to 7.4). The pipette solution consisted of 97.5 mM K-D-gluconate, 32.5 mM KCl, 0.5 M EGTA, and 10 mM HEPES (pH adjusted to 7.2). The membrane potential was held at -80 mV, and I-V relationships were generated using voltage steps from -70 to +70 mV in 10-mV increments.

For $I_{CRAC}$ measurements, the baseline bath solution contained 139 mM NaCl, 1 mM $MgCl_2$, 5mM KCl, 10 mM glucose, 1 mM EGTA, and 10 mM HEPES (pH 7.4). 1 µM thapsigargin and 1 µM tetrodotoxin were added to the bath prior to baseline recording. To eliminate cytosolic $Ca^{2+}$ and block $K^+$ currents, the pipette solution contained 145 mM Cs-methanesulfonate (Sigma, C1426), 8 mM NaCl, 10 mM $MgCl_2$, 10 mM EGTA, and 10 mM HEPES (pH adjusted to 7.2 with CsOH). To trigger $Ca^{2+}$ influx, an equal volume of $Ca^{2+}$-containing solution (140 mM NaCl, 4 mM $CsCl_2$, 2 mM $MgCl_2$, 10 mM $CaCl_2$, and 10 mM HEPES) was applied. Inward currents were measured at -80 mV every 5 seconds, with the membrane potential held at +30 mV between sweeps. Background noise, as defined by the software, was not subtracted during $I_{CRAC}$ calculations.

**ER-Tracker staining and classification**

HeLa cells seeded in 8-well chamber slides (Thermo Scientific, 155411) were stained with 1 µM ER-Tracker Green (Invitrogen, E34251) and 2 µg/mL Hoechst 33342 (Sigma, B2261) in ECB for 15 minutes at 37°C. Following staining, cells were washed and maintained in ECB for imaging. To distinguish morphologically distinct ER subdomains, images were analyzed using MATLAB. The ER silhouette was masked based on the ER-Tracker Green signal. Fluorescent intensity (ER signal) and edge gradient magnitude (ER difference) were calculated. Pixels with low gradient magnitudes were classified as sheet-like (lamellar) ER, whereas those with high gradient magnitudes were classified as tubule-like ER. The total number of pixels in each category was then quantified using MATLAB.

**Immunofluorescence (IF) staining**

For IF experiments, HeLa cells were seeded into 8-well chamber slides that had been pre-coated with 30 µg/mL collagen for at least 3 hours. On the day of the experiment, cells were rinsed with PBS and fixed with 4% paraformaldehyde in PBS for 10 minutes. Permeabilization was performed using 0.1% Triton X-100 for 10 minutes, followed by blocking in 5% BSA for 1 hour at room temperature. Cells were then incubated overnight at 4°C with primary antibodies, including anti-CLIMP-63 (Abcam, ab152154) and anti-FLAG (Cell Signaling, #8146). The following day, the wells were washed and incubated with Alexa Fluor-conjugated secondary antibodies (Invitrogen) and 300 nM DAPI (Invitrogen) for 1 hour at room temperature. After thorough washing, the slides were kept in PBS for subsequent imaging.

**Quantitative PCR (qPCR)**

Cells were lysed, and total RNA was isolated using NucleoZOL (Takara) according to the manufacturer's instructions. For reverse transcription, 4 µg of extracted RNA was transcribed using SuperScript™ IV Reverse Transcriptase (Invitrogen). Quantitative PCR was performed with SYBR Green Master Mix (Bio-Rad) using the Bio-Rad CFX Duet Real-Time PCR System. The specific primer sequences (5' to 3') used were as follows: *KCNT1* forward, 5’-CAGACCACCAGACCATCCTG-3', and reverse, TTGCACTCCTCCTCACACAC-3'; *STIM1* forward, 5’-GGACCTGTGGAAGGCATGGA-3', and reverse, 5’-CACTGAGCTGCAGCTTCCG-3'; *HPRT* forward, 5’-

AATTATGGACAGGACTGAACGTCTT-3', and reverse, 5’-ATGTAATCCAGCAGGTCAGCAA-3'. Data were collected and exported via Bio-Rad CFX Maestro Software, and subsequent analyses were performed using Python.

**Protein structural visualization and *in silico* analysis**

The predicted three-dimensional structure of human KCNT1 (isoform 2) was retrieved from the AlphaFold Protein Structure Database (accession number: AF-Q5JUK3-2-F1; version v6) [65-67]. Molecular visualization, mapping of charge distribution, and labeling of 740-DDE-742 and 1114-RRLSR-1118 were performed using the PyMOL Molecular Graphics System (Version 3.1.6.1, Schrödinger, LLC). Surface accessibility of the targeted charged motifs was assessed visually within the predicted structure.

**Multielectrode array (MEA) assay**

Transduced hiPSC-CMs were analyzed using the Axion Muse system. To confirm successful transduction prior to puromycin selection, mCherry expression was utilized as a fluorescent reporter. For FPD measurements, APs were paced via a single electrode applying a 5-mV stimulus for 125.75 ms, with a 300-ms interval between triggers. FPD data were exported using AxIS Analysis software. For spontaneous beating rate calculations, unpaced spontaneous APs were recorded. Waveforms were extracted using the Cardiac Data Plotting Tool and further analyzed in Python.
